# Bimodal genomic approach predicting Semaphorin 7A (SEMA7A) as prognostic biomarker in adrenocortical carcinoma

**DOI:** 10.1101/2025.04.03.647086

**Authors:** Anjali Dhall, Daiki Taniyama, Fathi Elloumi, Augustin Luna, Sudhir Varma, Suresh Kumar, Lauren Escobedo, Mirit I. Aladjem, Christophe E. Redon, Nitin Roper, William C. Reinhold, Jaydira Del Rivero, Yves Pommier

## Abstract

Adrenocortical carcinoma (ACC) is a rare and aggressive endocrine malignancy with high mortality and poor prognosis. To elucidate the genetic underpinnings of ACCs, we have analyzed the transcriptome data of 112 ACC tumor samples from patients enrolled in the TCGA and NCI. Among 72 bimodally expressed genes stratifying patients into prognostic groups, we focused on *SEMA7A*, as it encodes a glycosylphosphatidylinositol-anchored membrane glycoprotein (Semaphorin 7a) regulating integrin-mediated signaling, cell migration and immune responses. We find that high *SEMA7A* gene expression is associated with poor prognosis (hazard ratio = 4.27; p-value < 0.001). In hormone-producing ACCs, *SEMA7A* expression is elevated and positively correlated with genes driving steroidogenesis, aldosterone and cortisol synthesis, including *CYP17A1, CYP11A1, INHA, DLK1, NR5A1* and *MC2R*. Correlation analyses show that *SEMA7A* is co-expressed with the *integrin-β1, FAK* (focal adhesion kinase) and *MAPK/ERK* (mitogen-activated protein kinase/extracellular signal regulated kinases) signaling pathways. Immunohistochemistry (IHC) staining demonstrates the feasibility of evaluating *SEMA7A* in ACC tissues and shows significant correlation between gene expression (RNA-Seq) and protein expression (IHC). These findings suggest *SEMA7A* as a candidate for further research in ACC biology, a candidate for cancer therapy, as well as a potential prognosis biomarker for ACC patients.

**Translational relevance:** Adrenocortical cancer (ACC) remains a challenging disease primarily due to the scarcity of reliable biomarkers for predicting patient outcomes and informing innovative therapeutic strategies, as well as its rarity, which restricts the scope of clinical trials. In our study, we performed RNAseq and IHC analyses of ACC samples sourced from The Cancer Genome Atlas (TCGA), tissue microarray slide, and National Cancer Institute (NCI) cancer patient samples. Our findings indicate that a substantial proportion of ACC tumors exhibit expression of SEMA7A, a glycoprotein involved in Semaphorin cell surface signaling. Notably, elevated levels of SEMA7A were identified as a poor prognostic biomarker and were associated with activation of the integrin-ERK-MAPK kinase signaling pathways. These results suggest that ACC tumors with high SEMA7A expression should be considered at elevated risk, and SEMA7A may serve as a potential target for immunotherapeutic strategies, including antibody-drug conjugates, T-cell engagers, and/or small molecule inhibitors targeting the MAPK pathway.

## Introduction

Adrenocortical cancer (ACC) is a rare and aggressive malignancy arising from the adrenal cortex with high morbidity and mortality rates. The estimated annual incidence of ACC is ∼ 0.5 to 2 cases per million individuals worldwide. Notably, the incidence of ACC demonstrates a substantial age-related variation, with a peak occurring around the age of 50. In 40%–60% of patients, ACCs typically exhibit an aggressive biological behavior with symptoms of hormone hyperproduction (1). The prognosis for patients with ACC is generally poor, with a 5-year survival rate <15% among patients with distant metastases (2). Prognosis is closely linked to the stage at which the disease is diagnosed, with earlier-stage presentations associated with more favorable outcomes (3,4). The heterogeneous nature of ACC complicates its diagnosis and treatment, given the variability in clinical manifestations and prognostic outcomes (5). The precise molecular mechanisms underpinning ACC are not fully elucidated despite significant advancements in genomic and transcriptomic profiling. Scientific breakthroughs are warranted to identify diagnostic, prognostic, and therapeutic biomarkers, creating new opportunities for effective management strategies (6-9).

At present the clinical prognostic factors for ACC include tumor stage, cortisol secretion, and patient age. Tumor staging serves as the principal determinant of prognosis, as well as the status of surgical resection margins. Histological features such as mitotic activity exceeding 20%, a Ki-67 index greater than 12%, and a Weiss score greater than 6 have also been correlated with poor prognosis (10). Given the disease’s heterogeneity and rarity, investigations have focused on assessing various immunohistochemistry markers to identify more reliable prognostic factors. Efforts have also been made to develop genomic techniques for evaluating gene expression and alterations as potential molecular prognostic markers. Significant attention has been directed towards the Wnt/CTNNB1 and TP53 signaling pathways, which are frequently mutated in ACC, particularly with respect to CTNNB1 (beta-catenin) and P53 staining (11-13). The presence of somatic CTNNB1 mutations has been associated with recurrence in ACC patients (14).

Systemic therapies are typically employed in the adjuvant setting or for patients exhibiting metastatic or unresectable disease. Mitotane, an adrenolytic agent, has remained the most utilized medication for over fifty years. It is administered as adjuvant therapy following surgery or for inoperable or metastatic cases (15,16). In cases of advanced disease that are not suitable for surgical intervention, the utilization of cytotoxic agents in conjunction with mitotane is indicated.

Two commonly employed treatment regimens include the combination of etoposide, doxorubicin, and cisplatin with mitotane (EDP-M), and the combination of streptozotocin with mitotane (S-M). These therapeutic approaches were evaluated in an international phase-III clinical trial. The findings indicated that the EDP-M regimen was associated with higher objective response rates (ORRs) and improved progression-free survival (PFS) when compared to the S-M regimen; however, no statistically significant difference in overall survival was observed (17). Targeted therapies, including inhibitors of the insulin-like growth factor (IGF) pathway and immune checkpoint inhibitors, are currently under investigation in several clinical trials (18). Preliminary findings indicate that these treatments may provide new options for patients with advanced ACC, though further research is necessary to fully establish their efficacy and safety profiles.

To address the current challenges, research endeavors are directed towards our understanding of the genetic underpinnings of ACC (19-21). This step is crucial in the development of diagnostic and prognosis algorithms based on genomic data. Based on such approaches, several predictive characteristics have been identified, prognostic markers such as (BUB1B, PINK1, MKI67) and molecular classification derived from the clustering of genomic profiles, which proposed the presence of two primary transcriptional clusters, designated as C1A/C1B, exhibiting a strong correlation with survival outcomes. However, the C1A/C1B classifier faces challenges in its implementation as a prognostic biomarker owing to its inherent complexity (5,21-24).

Alternatively, molecular profiling approaches based on bimodal gene expression have been used to identify biomarkers possessing distinct expression distribution allowing for the classification of samples into two clearly defined expression states (25). For instance, estrogen receptor (*ESR1*) is a well-known bimodal expressed gene in breast cancer patients (26), and Schlafen 11 (*SLFN11*), a bimodal predictive genomic biomarker, predicts response to DNA-targeted chemotherapy (27). In this study, we implemented a statistical approach to identify bimodally expressed genes using transcriptome profiles of 112 ACC tumor samples from both the TCGA database and from patients enrolled in NCI clinical trials. We report that a cell surface marker, Semaphorin 7a (*SEMA7A)*, is significantly associated with poor prognosis, differentially up-regulated in ACC in comparison with normal adrenal tissues and reflects activation of the FAK and MAPK/ERK signaling pathways in a subset of ACC patients.

## Material and Methods

### Dataset Collection

In this study, we utilized transcriptomic and clinical data from TCGA-ACC and NCI-ACC database. For the NCI patient samples, a single institution study was conducted in accordance with recognized ethical guidelines as per The Belmont Reports and the Department of Health and Human Services Common Rule and was approved by the NIH Institutional Review Board. Written informed consent was obtained from all participants, and consent was approved by the NIH Institutional Review Board.

RNA-seq sequencing data processed using NCI CCBR RNA-seq pipeline (https://github.com/skchronicles/RNA-seek.git) and STAR (2.7.11b) to align reads to the hg38 reference genome. RSEM was used to normalize gene expression values expressed as log2(FPKM+1) and “RemoveBatchEffect” function from the “Limma” package used to remove the impact of the library preparation protocols. In total, 112 ACC patients from TCGA-ACC (n=79) and NCI-ACC (n=33) were included in this study (see Table 1, Table S1). Additionally, normalized RNA-seq expression data from multiple TCGA cancer types (including BLCA, BRCA, CESC, CHOL, ESCA, HNSC, KIRC, KIRP, LAML, LIHC, LUAD, LUSC, PAAD, PRAD, THCA, UCS, and UVM) were obtained from cBioPortal and normal tissue expression datasets from Genotype-Tissue Expression (GTEx) database.

**Table 1:** Clinical and demographic characteristics of ACC patients.

| **Variable** | **Number of Patients**<br>**(total = 112)** |
| --- | --- |
| **Age** |  |
| Median age < 49 years | 56 |
| Median age ≥ 49 years | 56 |
| **Gender** |  |
| Female | 77 |
| Male | 35 |
| **Tumor Stage** |  |
| Stage I | 12 |
| Stage II | 50 |
| Stage III | 18 |
| Stage IV | 29 |
| Not Available | 3 |
| **Vital Status** |  |
| Alive | 75 |
| Dead | 37 |
| **Median overall Survival (OS)** |  |
| < 43 months | 58 |
| ≥ 43 months | 54 |
| **Hormone Producing** |  |
| Yes | 64 |
| No | 35 |
| Not Available | 13 |

### Bimodal Approach

To identify RNA-seq expression bimodality we employed a combination of the Hartigan Dip test (using the R *Diptest* package, R version 4.2.3) and Normal/Gaussian Mixture Models (with the R *nor1mix* package using an Expectation-Maximization (EM) approach). For each individual gene, the expression distribution was sorted and tested for bimodality using the Dip test. If the distribution was found bimodal using Dip test with p-value <0.05, we applied the *norMixEM* function from the R *nor1Mix* package to fit a mixture of two normal distributions. Once the fitting had converged, the parameters of this model, such as the means (*µ*_*1*_, *µ*_*2*_), weights (*w*_*1*_, *w*_*2*_ with *w*_*1*_+ *w*_*2*_=1), and standard deviation (*σ*_*1*_, *σ*_*2*_) for the two distributions, along with the log-likelihood, were computed per gene. We calculated an overall bimodal score, as a rough analog of a t-statistic, as the absolute difference in means divided by the larger of the two standard deviations (See Equation 1). An additional criterion was that the difference between the weights for the two distributions was less than 0.25 to ensure the selection of genes with similar number of samples in the high and low expression distributions.

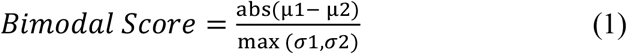

### Statistical Analyses

Univariate survival analyses were performed using the Cox proportional hazards (Cox-PH) regression model. Gene transcripts from ACC samples were stratified into two categories high and low expression based on median expression cut-off. Significant differences in survival distributions between the high-risk (high expression) and low-risk (low expression) groups were assessed using the log-rank test, with the results presented in terms of Hazard Ratios (HR), p-values, confidence interval and concordance index. HR >1 indicates a negative impact on patient survival, while HR <1 improved survival. An HR of 1 indicates no effect on survival. The stratification of patients into high-risk and low-risk groups was visually represented using Kaplan-Meier (KM) survival curves, providing a distinct depiction of the survival probabilities over time for each group. We used the “survival” and “survminer” packages in R (version 4.2.3). Correlation analyses were performed using the Pearson correlation test. Mann–Whitney U test analyses were performed using GraphPad Prism 9.0 software (GraphPad Software Inc.).

### Tissue Microarray and Immunohistochemistry (IHC) Evaluations

Tissue microarray slide AG991 was used to stain ACC and normal adrenal gland tissue samples. Immunohistochemistry (IHC) staining was performed on LeicaBiosystems’ BondRX autostainer with the following conditions: Epitope Retrieval 2 (EDTA) 20’, SEMA7A (Santa Cruz #sc-374432, 1:100 incubated 30’), and the Bond Polymer Refine Detection Kit (LeicaBiosystems #DS9800). Isotype control reagent (mouse IgG2a, BD Biosciences #553454) was used in place of primary antibody for the negative control. Slides were removed from the Bond autostainer, dehydrated through ethanol, cleared with xylene, and coverslipped. Slides were scanned using a Leica Aperio AT2 scanner (Leica Biosystems, Buffalo Grove, Illinois) at 20X magnification. The stained IHC slides were quantified using the HALO image analysis software 3.6 (Indica Labs, Albuquerque, NM, USA). Automated quantification of percent positive cells for each core was performed using cytonuclear algorithm version 2.0.5.

For the NCI-ACC patient specimens, formalin-fixed paraffin-embedded tissue sections were used for IHC. SEMA7A IHC was performed using a previously validated method (28). Optimized staining protocol included antigen retrieval performed by microwave heating in Antigen Retrieval Buffer (pH 6.0) (#ab93678, Abcam) for 20 min. The sections were incubated with mouse monoclonal anti-SEMA7A antibody (Santa Cruz #sc-374432, 1:100 dilution in TBST (1×TBS + 0.5% Tween 20)) overnight at 4°. The sections were then incubated for 30 minutes in N-histofine Simple Stain MAX PO(MULTI) (Nichirei Biosciences Inc., # NIC-414151F) followed by 5 minutes incubation with chromogen DAB-3S (original dilution by manufacturer, Nichirei Biosciences Inc.; #415192F) at room temperature.

## Results

### Overall Workflow and Implementation of the Bimodal Approach

The overall workflow for the selection of bimodal genes is provided in Figure 1A. At first, we applied the bimodal algorithm to the gene expression data collected from both TCGA and NCI. Ranking the genes based on their bimodal scores led to the selection of 200 genes having the highest bimodal distribution. Analysis on those top 200 genes retrieved 72 genes significantly associated with the survival of ACC patients. Among them 54 are significantly associated with poor prognosis, and 18 with better overall survival (Figure 1A). The list of the 72 bimodal-survival genes is provided in (Table S2) along with their annotation as belonging to the C1A/C1B molecular subtypes. We also assigned the NCI-ACC patients as C1A/C1B molecular subtypes using the ConsensusClusterPlus approach previously used for the TCGA-ACC samples (19,29) (Figure 1B, upper section labeled as molecular subtypes in red for C1A and blue for C1B).

**Figure 1:**
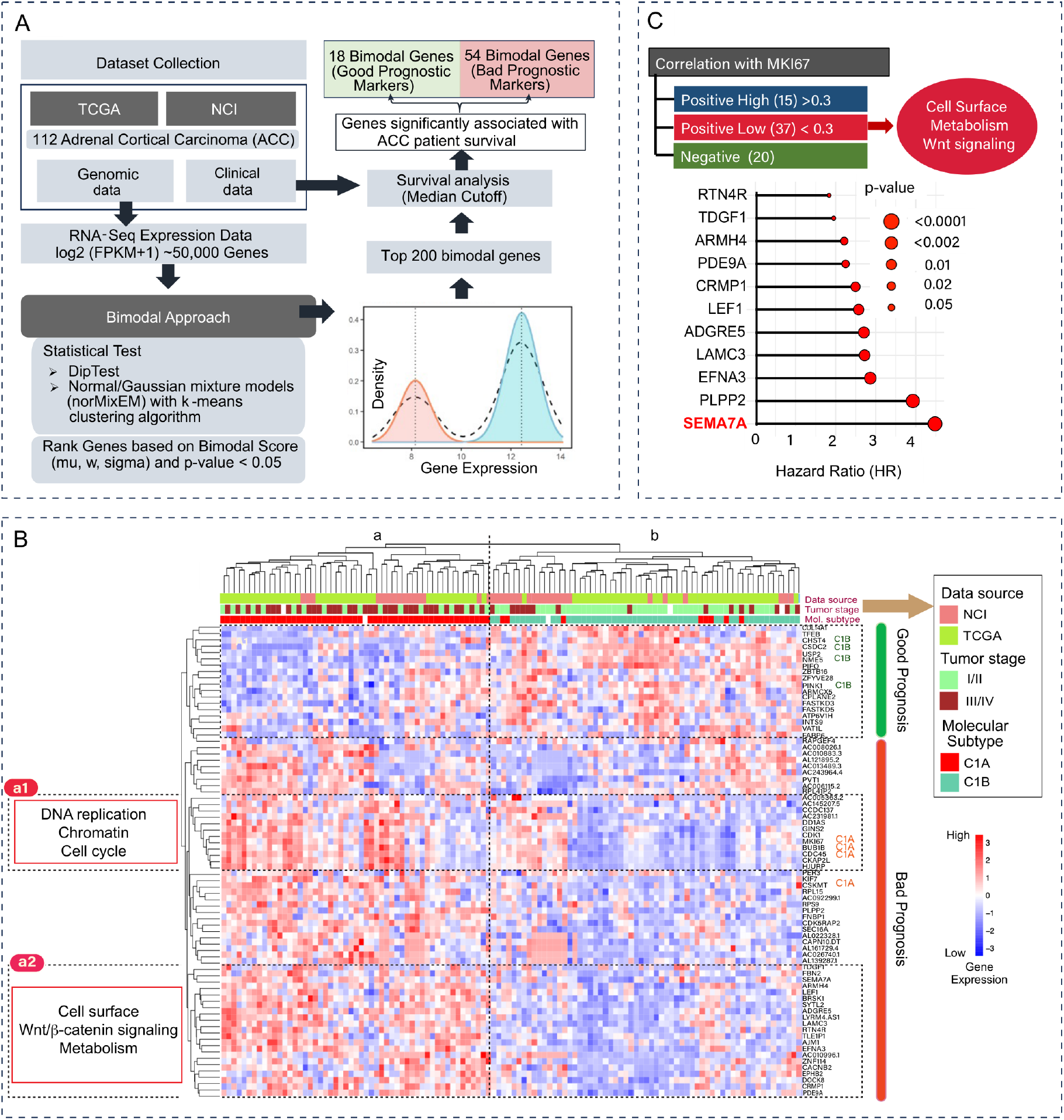
(**A**) Study architecture and bimodal gene selection. (**B**) Unsupervised hierarchical clustering heatmap displaying 72 bimodal genes associated with ACC patient survival, tumor stage, molecular subtype (C1A/C1B), and *PINK1, BUB1B, MKI67* signature. (**C**) Classification of the 72 bimodal genes into 3 groups based on correlation analysis with *MKI67*. Subgroup **a1** is enriched for DNA replication markers and subgroup **a2** for cell surface markers including *SEMA7A*.

Next, we computed a double clustering heatmap displaying both patient samples (columns in Figure 1B) and expression of the 72 bimodal-survival genes (rows). Patient distribution formed two main clusters (labeled “a” and “b” in Figure 1B). Cluster “**a**” is enriched for patient with advanced tumors (stage III & IV) and C1A subtype while it is the reverse for patients in cluster “**b**” (high fraction of localized tumors with C1B staging). Analysis of the gene distribution (rows in Figure 1B) shows an upper cluster encompassing genes that tend to be highly expressed in the cluster “**b**” patients and can be considered good prognosis genes. Consistently, among them, 4 are known C1B genes: *CHST4* (encoding CarboHydrate sulfotransferase 4), *CSDC2* (encoding Cold Shock Domain Containing C2), *NME5* (encoding Non-Metastasis cell 5 protein) and *PINK1* (encoding PTEN-Induced Kinase 1, whose high expression is a well-established good prognosis biomarker in ACC (21,29). The bimodal-survival genes associated with poor prognosis (cluster “a” in Figure 1B), are distributed in 4 clusters. Two of them annotated as “a1” and “a2” in Figure 1B. Cluster “**a1**” include genes associated with DNA replication, chromatin and cell cycle: *HJURP, CDK1, CDCA5, GINS2, DDIAS*. Cluster “**a2**” includes cell surface markers, metabolism and Wnt-signaling-related genes including *PDE9A, BRSK1, SYTL2, RTN4R, EFNA3, PLPP2, CRMP1, LAMC3, LEF1* and *SEMA7A*.

Further analyses were performed to divide the bimodal genes based on their relationship with *MKI67* and identify novel predictive biomarkers that would complement *MKI67*. Thirty-four genes showed a low correlation with *MKI67* (Figure 1C). Among them *SEMA7A*, which encodes the cell surface marker Semaphorin 7A showed the highest Hazard Ratio (HR = 4.27 and significant p-value < 0.001) (Figure 1C). These results reveal SEMA7A expression as a novel prognosis biomarker with a broad bimodal expression in ACC.

### High expression of SEMA7A in ACC

In the TCGA-pan cancer datasets, ACC is with pancreatic adenocarcinoma (PAAD) and bladder cancer (BLCA) among the cancers expressing the highest levels of *SEMA7A* (Figure 2A). This high expression is specific to ACC cancer patients as the expression of *SEMA7A* is low in normal adrenal gland (Figures 2B and S1). The normal tissues expressing highest levels of *SEMA7A* are lymphoid, nervous and germinal (Figure S1). We next examined whether *SEMA7A* expression is associated with clinical stage, hormone production and gender. Indeed, we observed that highest expression of SEMA7A was observed in high-grade (III/IV) and hormone-producing ACC (Figure 2C). Female patients tended to have higher SEMA7A expression than males, but the difference was not significant (Figure 2C). The prognosis value of SEMA7A is shown in Figure 2D with significantly reduced survival of patients with high *SEMA7A* (hazard ration (HR) = 4.27 and p-value <0.001). As expected, stages III/IV and hormone-producing tumors are also at significantly higher risk (Figure 2E). Not significant difference was observed between males and female patients (Figure 2E). Correlation analyses showed a significant association of *SEMA7A* expression with the expression of adrenocorticotropic hormone receptor, steroidogenic enzymes, cholesterol transporters and their transcriptional regulator genes (*CYP11A1, CYP17A1, MC2R, NR5A1/SF1, DLK1* and *INHA*) (30-35) (Figure 2F).

**Figure 2:**
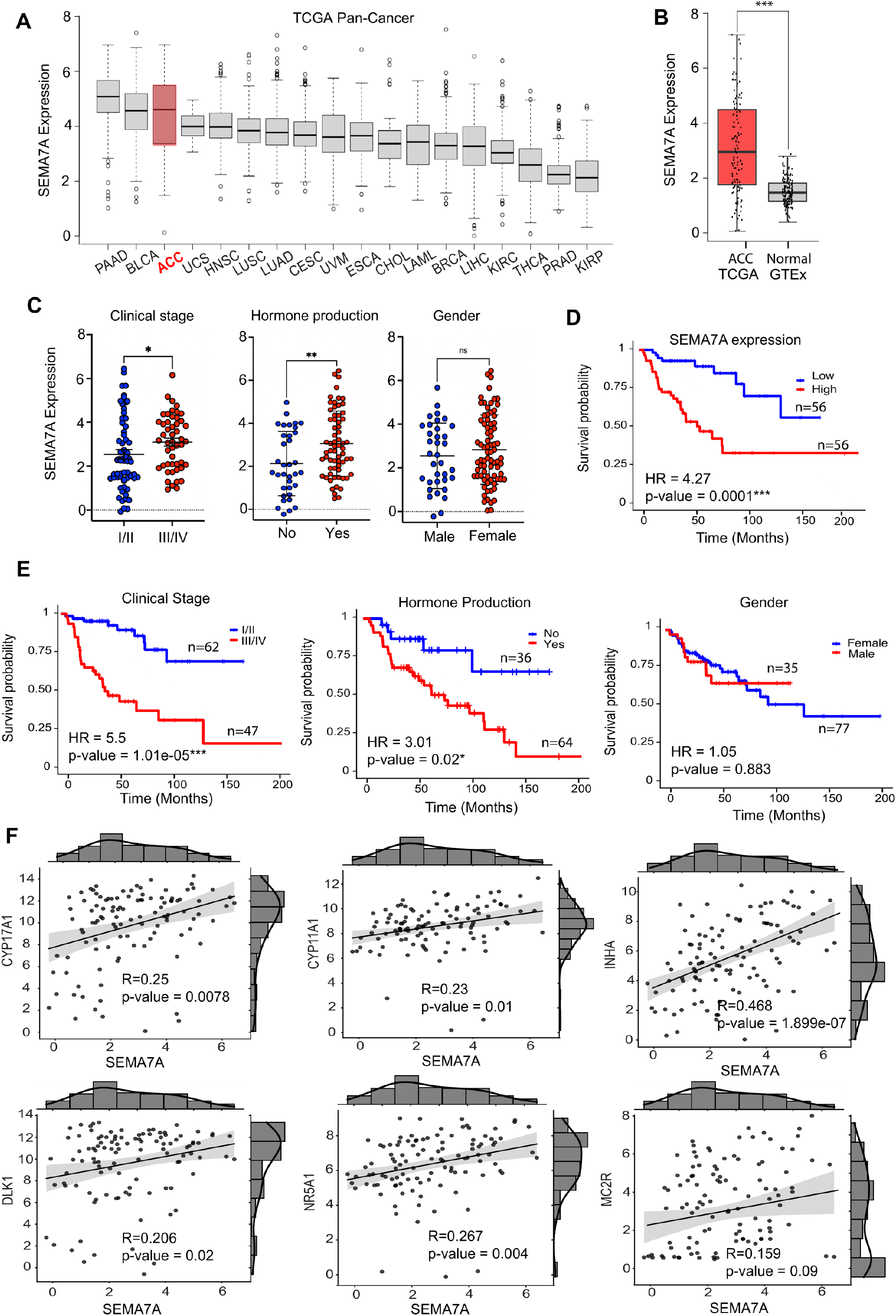
Expression of *SEMA7A* in ACC and SEMA7A expression as prognosis biomarker in ACC. **(A)** *SEMA7A* expression in various TCGA cancer types. **(B)** Differential expression and bimodal distribution of *SEMA7A* in normal adrenal gland (GTEx) and ACC. **(C)** Expression of *SEMA7A* as a function of tumor stage (III/IV vs. stage I/II), hormone production and gender of ACC patients. **(D, E)** Kaplan–Meier survival plots demonstrating the significant association of *SEMA7A* expression with clinical stage and hormone production in ACC patients.

### Activation of the SEMA7A-integrin-β1 downstream signaling pathways in ACC

SEMA7A is known to belong to a SEMA7A-integrin-β1 axis that activates Mitogen-Activated Protein Kinase (MAPK) cascades; especially the ERK/MAPK signaling pathway, which plays crucial role in regulating oncogenic functions, including metastasis and tumor progression (36-39). As shown in Figure 3A, expression of the downstream pathway genes *ITGB1, AKT, PTK/FAK, ERK/MAPK1, LIMK1, CFL1* shows significant positive correlation with *SEMA7A* expression and with each other. These observations are consistent with the functional activation of the downstream pathways of SEMA7A in the ACC tumors expressing high *SEMA7A* transcripts.

**Figure 3:**
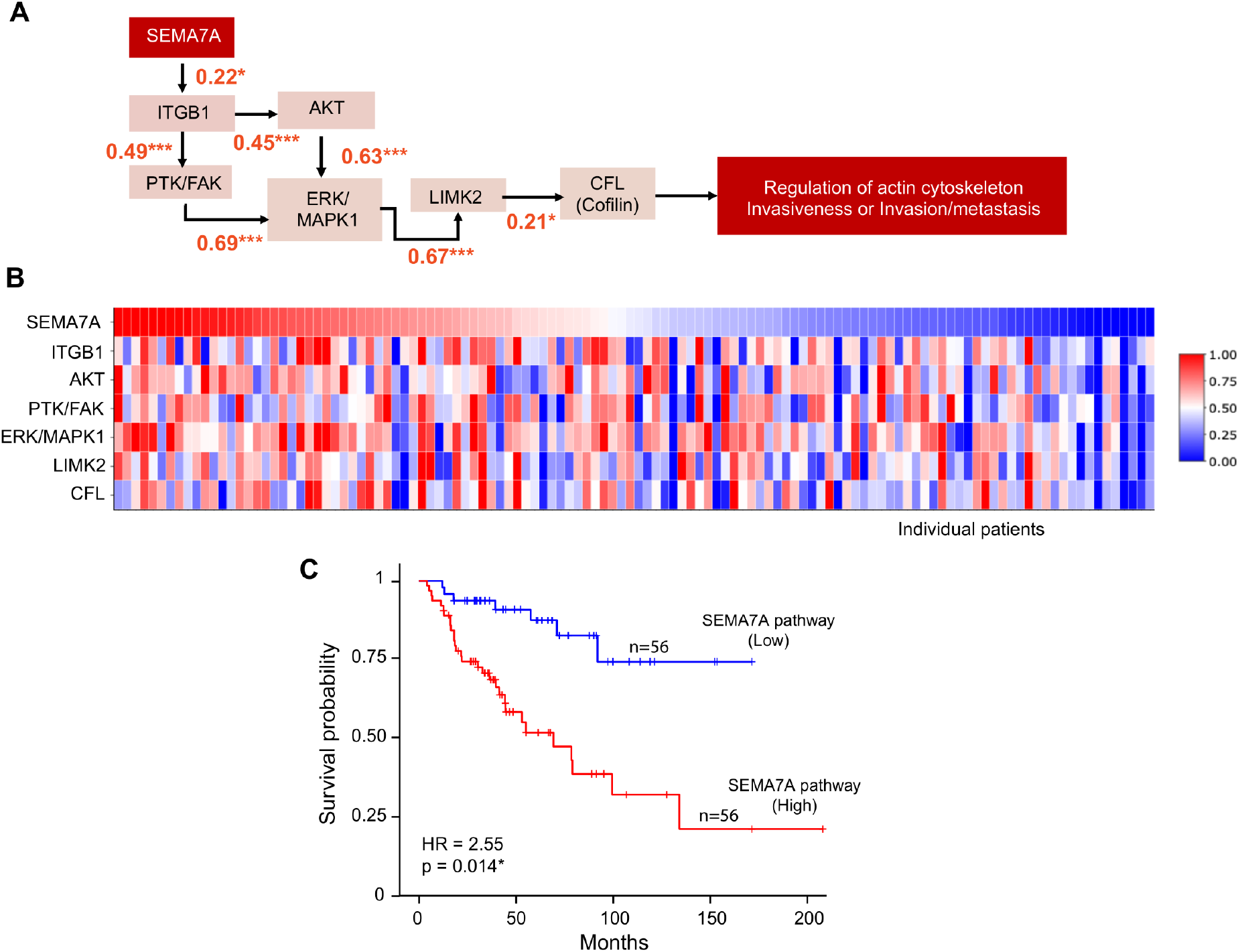
**(A)** MAPK/ERK signaling pathway is co-regulated via SEMA7A, **(B)** Heatmap shows the expression of neighboring genes of MAPK/ERK pathway in ACC patients. **(C)** Kaplan– Meier survival plot, based on average expression of SEMA7A pathway genes, divided into two groups based on median expression cut-off.

Expression heatmap of SEMA7A-integrin-β1 downstream pathway genes stratifies the ACC patient samples into two subgroups, one having high expression and the other having low expression of those genes (Figure 3B). To explore the association between the downstream genes of the SEMA7A-integrin-β1 pathway and overall survival in ACC patients, we calculated the average expression of genes involved in the SEMA7A pathway and stratified the patients into two groups based on the median expression value (cut-off). The survival analysis shows the poor prognosis of patients having high average expression of SEMA7A pathway genes (Figure 3C).

Together these results demonstrate the functionality of the SEMA7A pathway in a significant fraction of ACC tumors and the significantly poor prognosis of patients with high *SEMA7A* expression and SEMA7A pathway activation.

### Exploration of SEMA7A protein expression in ACC

To compare SEMA7A transcripts and protein expression, we first used tissue microarray slides to determine whether SEMA7A protein expression could be measured in normal adrenal gland (n=6), adrenocortical adenoma (n=27) and ACC (n=14) tumor tissues. Figure 4A shows representative immunohistochemistry (IHC) staining images demonstrating the fraction of cells expressing SEMA7A in ACC tissues compared to normal adrenal gland tissue and adrenocortical adenoma (benign tumor). Quantification (Figure 4B) is consistent with the bimodal distribution of SEMA7A expression observed at the transcript level. Indeed, some samples show very high percentage of positive cells expressing SEMA7A while other samples display low fractions. This suggests that SEMA7A protein levels determined by IHC can potentially be used as a biomarker to identify specific subgroups of ACC patients and score tumor aggressiveness instead of or in conjunction with RNA-Seq analyses.

**Figure 4:**
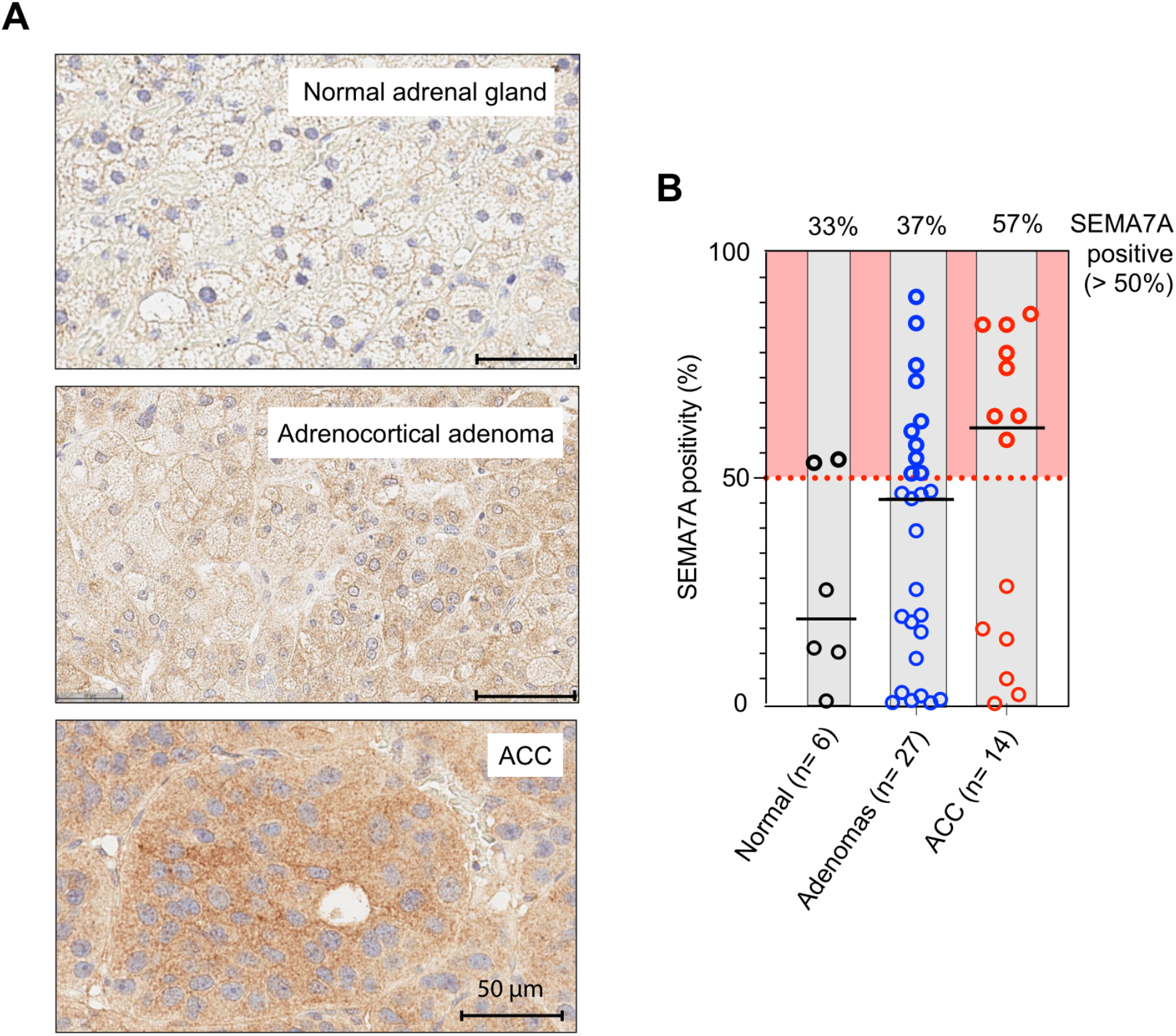
**(A)** Immunohistochemistry (IHC) examination of SEMA7A protein expression in normal adrenal gland tissues, adrenocortical adenomas (benign) and adrenocortical carcinomas (ACC). **(B)** Percentage of SEMA7A positive cases in normal, benign and malignant tissue samples. Note the bimodal distribution of SEMA7A in ACC.

To validate the IHC approach in ACC tumor samples, we compared the protein and RNA expression levels of SEMA7A in the NCI-ACC samples. We selected three samples having low and three having high SEMA7A RNA-Seq expression levels. As shown in Figure 5, the samples having low RNA-Seq levels do not show high expression at the protein level. By contrast, patients with high RNA-seq expression of SEMA7A (≥ 5) show protein expression. These results indicate a positive correlation between RNA-Seq expression and protein levels in the selected ACC samples and suggest the value of measuring SEMA7A both by RNA-Seq and IHC to classify ACC patients.

**Figure 5:**
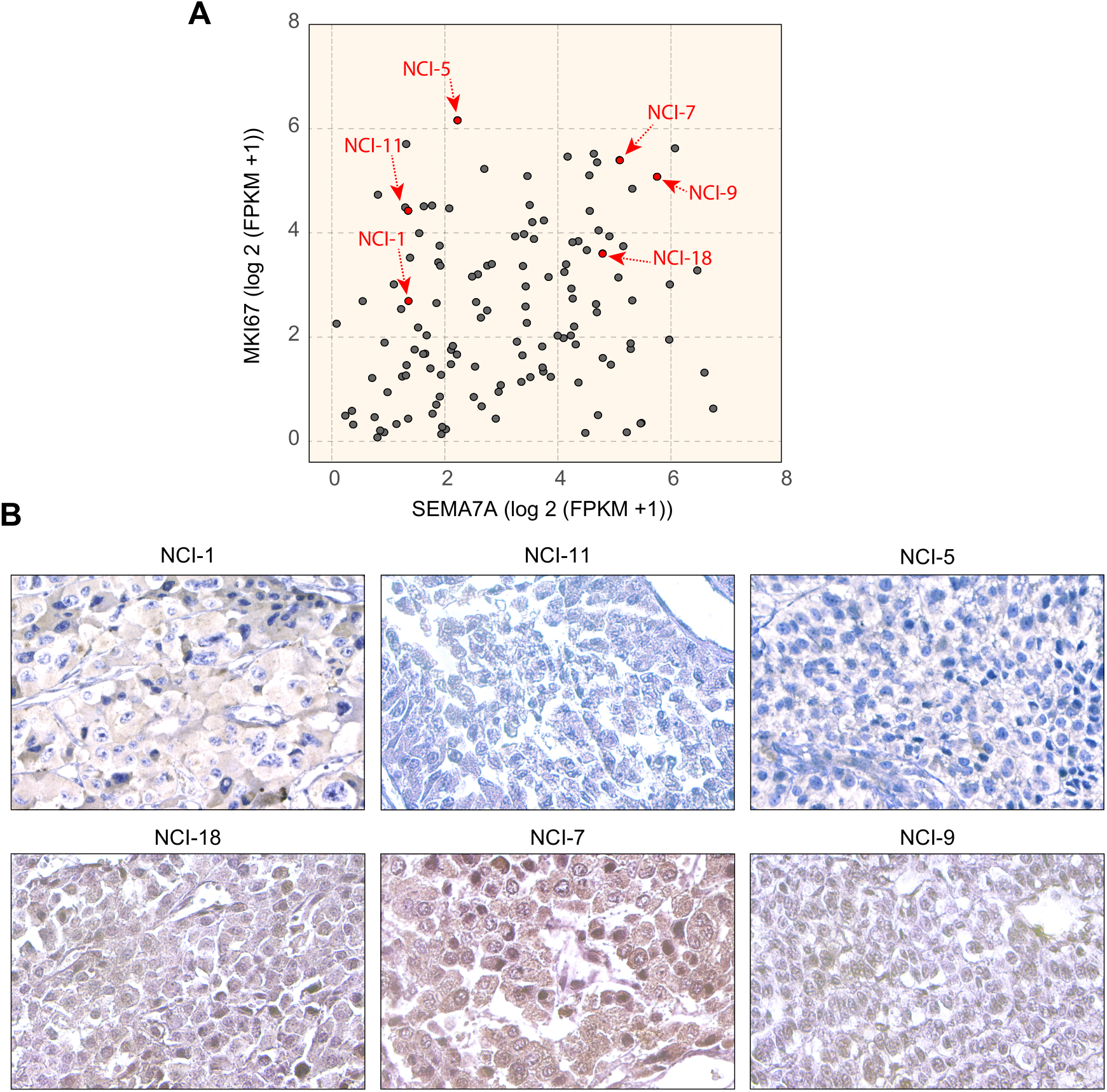
**(A)** Wide distribution of SEMA7A expression measured by RNA-Seq in 112 ACC tumor samples (see Figure **1A**) and lack of correlation with MKI67 expression. **(B)** Representative immunohistochemistry (IHC) staining of SEMA7A in three samples with high vs. low expression of SEMA7A in NCI-ACC patients.

## Discussion

In this study, we have implemented a bimodal approach using Dip test and Normal/Gaussian Mixture Models that led to the identification of SEMA7A as one of the genes with a broad range of bimodal distribution in ACC. Previously, Justino et al. introduced an integrated method to identify bimodal genes across multiple TCGA datasets (40). However, their analysis did not include ACC samples (26,40,41). Our aim was to explore a novel approach to classify ACC patients and capture heterogeneity across ACC tumors.

ACC presents considerable challenges due to its rarity, the complexity of its biology and its poor prognosis. Furthermore, due to the restricted number of cell lines and preclinical models, ACC patients have limited treatment options. Surgical resection continues to be the foremost approach for the treatment of localized cases, while metastasectomy has been proposed for the management of recurrent or metastatic ACC. This strategy carries significant therapeutic and palliative implications and is only associated with prolonged survival in carefully selected patients. Due to the prevalence of surgical interventions, high quality samples for biological and genomic analyses are routinely available at the Center for Cancer Research (CCR) of the National Cancer Institute at the National Institutes of Health (see Figs. 1 & 5).

To identify novel biomarkers for ACC, we have developed a method to identify bimodally expressed genes in ACC patient tumors to stratify patients based on their overall survival and discover predictive biomarkers. At first, we used RNA-seq expression profiles of ACC patient tumors samples and computed the bimodal score and p-value for each gene. Our analysis revealed 72 bimodal genes having significant association with the overall survival of ACC patients. SEMA7A came up as the gene with the most significant association with prognosis independently of *MKI67* (see Fig. 5A). In our analysis, we have observed that *SEMA7A* is significantly up regulated in ACC as compared to normal adrenal gland tissues (see Fig. 2). *SEMA7A* belongs to the Semaphorin gene family. It encodes SEMA7A (CDw108), a signaling glycoprotein (as its name indicates) anchored to the cell surface via glycosylphosphatidylinositol (GPI) linkage (42). Upon cleavage of the GPI membrane anchor, SEMA7A exists both as shed or membrane-bound. SEMA7A influences cellular dynamics in both cell-autonomous and non-autonomous manners. It enhances the mobility of melanocytes, neuronal cells and immune cells through the activation of β-1 integrin signaling pathways (37,38,43). In its normal physiological role, SEMA7A binds β1-integrin to activate downstream signaling cascades, including the pro-invasive MAPK/ERK and pro-survival PI3K/AKT pathways (44).

Recent findings have identified SEMA7A as a tumor promoter, accelerating tumor growth, increasing invasiveness, promoting adhesion to tumor-associated extracellular matrices. SEMA7A induces epithelial to mesenchymal transition, promoting metastasis in melanoma, glioblastoma, oral, kidney and breast cancers (36,45-49). Moreover, high expression of SEMA7A has been associated with significantly decreased patient survival in ER+ breast cancer patients (50). Prior to this work, no association between SEMA7A and ACC have been observed. Relevant to our study, in neuronal cells of the hypothalamus, SEMA7A expression is regulated by steroid hormones (51); an association we found also in ACC (see Fig. 2). SEMA7A is also upregulated in high grade (Stage III/IV) and hormone producing ACC patients (see Fig. 2). Our correlation analyses show that the SEMA7A-integrin-β1 axes activates the ERK/MAPK signaling pathway in the subset of ACC patients with poor prognosis (see Fig. 3).

Immunohistochemistry (IHC) staining of ACC tissues shows bimodal distribution of SEMA7A in ACC patients with lower levels in normal adrenal tissues and benign adrenal tumors (see Figs. 4 & 5). The positive correlation between RNA-seq and immunohistochemistry (IHC) of NCI-ACC patient samples suggests that SEMA7A could be a promising prognostic biomarker for clinical applications. SEMA7A as a potential biomarker for further investigations and for application in clinical diagnostics, particularly for forecasting outcomes in ACC patients. Finally, SEMA7A expression on the ACC tumors position it as a potential target for the development of immunotherapies, including antibody drug conjugates (ADC) and T-cell engagers.

## Acknowledgements

Our work is supported by Center for Cancer Research, the Intramural Program of the National Cancer Institute, NIH (Z01-BC 006150). We wish to thank Jonathan Hernandez, CCR Surgical Oncology Program for the surgical samples.

## Author Contribution

Y.P. supervised the study. Y.P., A.D and J.D.R. devised the concept. Y.P., A.D., F.I., A.L., S.V., W.C.R. and N.R. performed data analysis. D.T., S.K., C. E. and L.E., performed immunohistochemistry staining and analysis. A.D., J.D.R., Y.P., M.I.A and S.V. wrote and critically reviewed the manuscript. All authors reviewed the results and approved the final version of the manuscript.

## Supplementary Data

**Figure S1:**
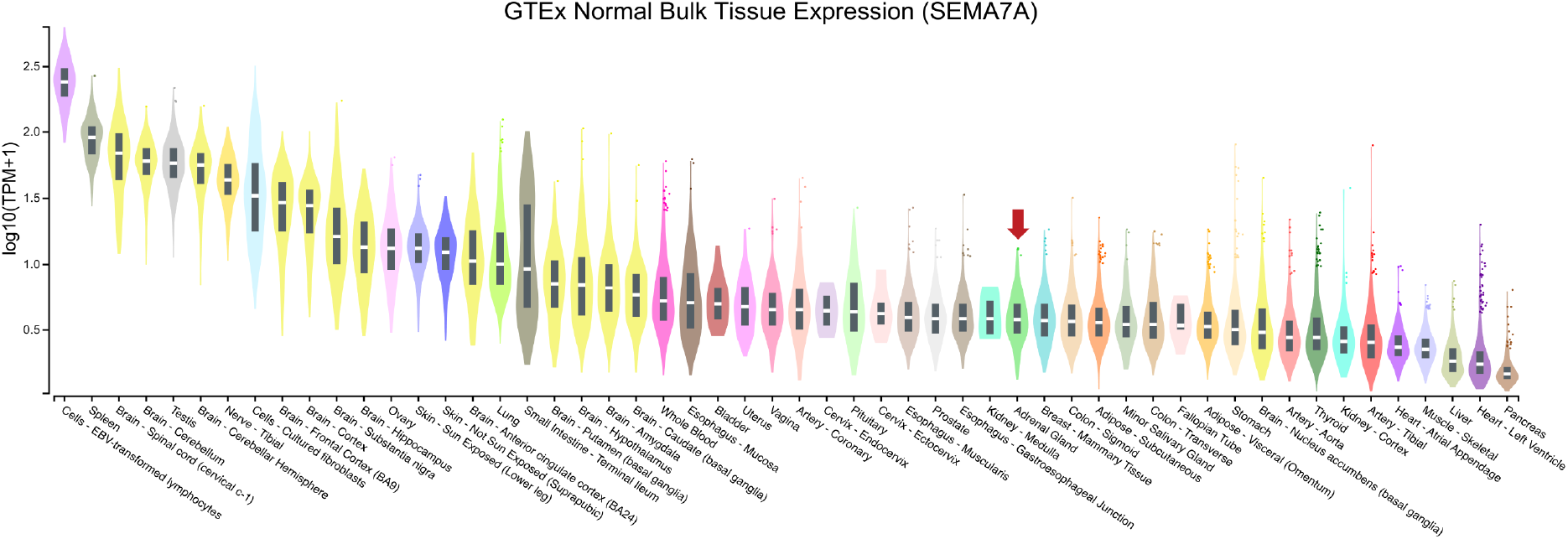
Distribution of SEMA7A expression across a wide range of normal human tissues obtained from GTEx database.

